# Lesion network mapping of depressive rumination identifies candidate brain stimulation targets

**DOI:** 10.64898/2026.11.23.768538

**Authors:** Zhen Fan, Yu-Yang Chen, Hai-Juan Li, Taicheng Huang, Di Yuan, Xiao-Yu Wang, Xiao-Na Zhu, Yu Zhang, Zheng-Wei Yuan, Wen-Jie Zhou, Chao-Gan Yan, Ti-Fei Yuan, Xiao Chen

## Abstract

Rumination is a recurrent dwelling on depressed feelings and a prominent feature of depression and other mood disorders, yet its neural organization remains incompletely understood. Here we integrate lesion network mapping, functional MRI, and intracranial electrophysiology across five datasets to delineate a brain circuit associated with rumination. In patients with diffuse glioma (*n* = 59), functional connectivity (FC) from lesions was estimated from a normative connectome database (*n* = 3001) and associated with individual ruminative traits. The resulting map was highly consistent across connectomes derived from individuals with depression and healthy controls. Its spatial organization followed the cortical modulation–representation gradient and showed associations with synaptic gene-expression profiles and neurotransmitter receptor distributions. Testing against random-coordinate null models identified anterior and posterior ventrolateral prefrontal and temporoparietal nodes. Independent task datasets provided convergent functional evidence for the anterior ventrolateral prefrontal cortex (aVLPFC) and temporoparietal junction (TPJ). Induced rumination increased local functional synchrony (ReHo) in the identified significant rumination nodes and their functional coupling with the frontoparietal network (FPN). Intracranial recordings revealed reduced gamma band power in aVLPFC and increased delta band power in TPJ during rumination. These findings link lesion-associated variation in rumination to distinct local neural dynamics and distributed functional interactions. They provide a lesion-informed framework for investigating persistent negative thought and nominate candidate sites for prospective neuromodulation studies.

## Introduction

Rumination in major depressive disorder (MDD) and other psychiatric conditions refers to automatic and persistent focus on depressed mood and its causes and consequences ^1,2^. It is a transdiagnostic behavioral element that plays a critical role in a number of common neuropsychiatric disorders, including MDD, obsessive-compulsive disorder (OCD), post-traumatic stress disorder, anxiety, etc. ^3^ Rumination is most strongly related to MDD, predicting the onset, severity, and duration of depression ^4^. It also remains elevated after remission from a depressive episode, increasing the likelihood of relapse ^5^. Thus, understanding the neural mechanisms of rumination is crucial for elucidating the pathophysiology and phenomenology of MDD and may inform the development of novel neuromodulation therapies for depression.

Existing neuroimaging studies have uncovered a brain circuit that is associated with rumination ^6^. Previous studies showed that rumination involves not only local activity, but also complex functional coupling among a wide range of brain regions that belong to several large-scale brain networks, including the default mode network (DMN), fronto-parietal network (FPN), and ventral and dorsal attention network (VAN and DAN) ^7,8^. In task fMRI studies using the rumination inducement task (RIT) ^9^, participants showed elevated brain activity in DMN regions such as the medial prefrontal cortex (MPFC), precuneus, and temporal parietal junction (TPJ) ^10^. Resting-state fMRI (rs-fMRI) studies found that the ruminative tendency measured with the ruminative response scale (RRS) ^11^ was associated with functional connectivities (FC) among DMN and FPN regions ^12,13^. We previously developed a rumination state task (RST) ^14^ that induces sustained, active rumination and used it to characterize the functional connectome of rumination. We identified static and dynamic patterns of functional coupling that primarily involve the DMN, FPN, DAN, and VAN ^14–17^. We also identified distinct alterations in intracranial EEG oscillatory activity during active rumination ^18^. Together, these findings suggest that rumination is supported by coordinated interactions among multiple higher-order brain networks.

While these brain networks have been consistently implicated in rumination, these approaches are inherently correlational, which limits their ability to translate into therapeutic targets ^19^. Brain lesions offer more direct evidence of causal relationships between symptoms and human neuroanatomy ^20^. Recently, brain lesion data along with the lesion network mapping (LNM) framework have been used to identify brain circuits underlying MDD ^21–24^, OCD ^25^, anxiety ^26^, psychosis ^27^, and nicotine addiction ^28^, and its clinical relevance has been initially supported by a randomized head-to-head transcranial magnetic stimulation (TMS) trial ^29^, whose targets were inspired by LNM ^30^. Accordingly, extending this approach to rumination would further validate the LNM framework and uncover novel stimulation targets for this symptom. Furthermore, the rumination circuit generated with LNM might provide additional information regarding the multi-scale neurobiological underpinnings of rumination, uncovering its relevance to the brain’s critical functional and structural architecture, such as functional gradients ^31,32^, gene expression ^33^, neurotransmitter receptors ^34^, and cell types ^35^.

In this study, we analyzed the association of lesion location with rumination across five independent datasets. Given recent methodological concerns ^36^ regarding LNM, we used a permutation-based approach ^37^ to construct an empirical null distribution for assessing the statistical significance of our findings and further validated the results using three independent datasets. We tested whether lesion locations that affect rumination connect to the same brain circuit, whether this circuit corresponds to a specific multi-scale neurobiological profile, and whether it could inspire novel potential neuromodulation targets for rumination.

## Results

### Lesions converge on a common rumination circuit

Using the GLI-RUM (*n* = 59) and DIRECT (*n* = 3001) datasets (Table 1), the rumination circuit map was generated based on the normative connectivity of lesions that influence ruminative tendency (Figure 1). The tumors were spatially diverse. Most tumors were located in the anterior cingulate cortex (ACC), insula, and dorsal medial prefrontal cortex (DMPFC). A minority of tumors were located in the visual cortex and motor cortex (Figure S1). We found a specific rumination circuit (Figure 2), which could be reproduced in two normative connectomes (MDD patients and HCs, Spearman’s rho = 0.997, *P* < 0.001). The global signal regression (GSR) did not alter this spatial pattern (Spearman’s rho = 0.949, *P* < 0.001). As a result, we adopted the lesion network map derived from an MDD-based normative connectome with GSR applied during preprocessing as our primary result. The rumination circuit was similar to the circuits of its two established subtypes (brooding: Spearman’s rho = 0.978, *P* < 0.001; reflection: Spearman’s rho = 0.965, *P* < 0.001). Rumination is a psychopathological process that is highly associated with depression. Accordingly, we found that the rumination circuit showed a significant yet weaker spatial correlation with the BDI-generated circuit (Spearman’s rho = 0.804, *P* < 0.001). Furthermore, the BDI circuit showed significant spatial correlation with the depression circuit developed by Siddiqi et al. ^30^ (Spearman’s rho = 0.313, *P* = 0.012), providing convergent evidence supporting the methodological validity of our approach (Figure S2).

**Figure 1.**
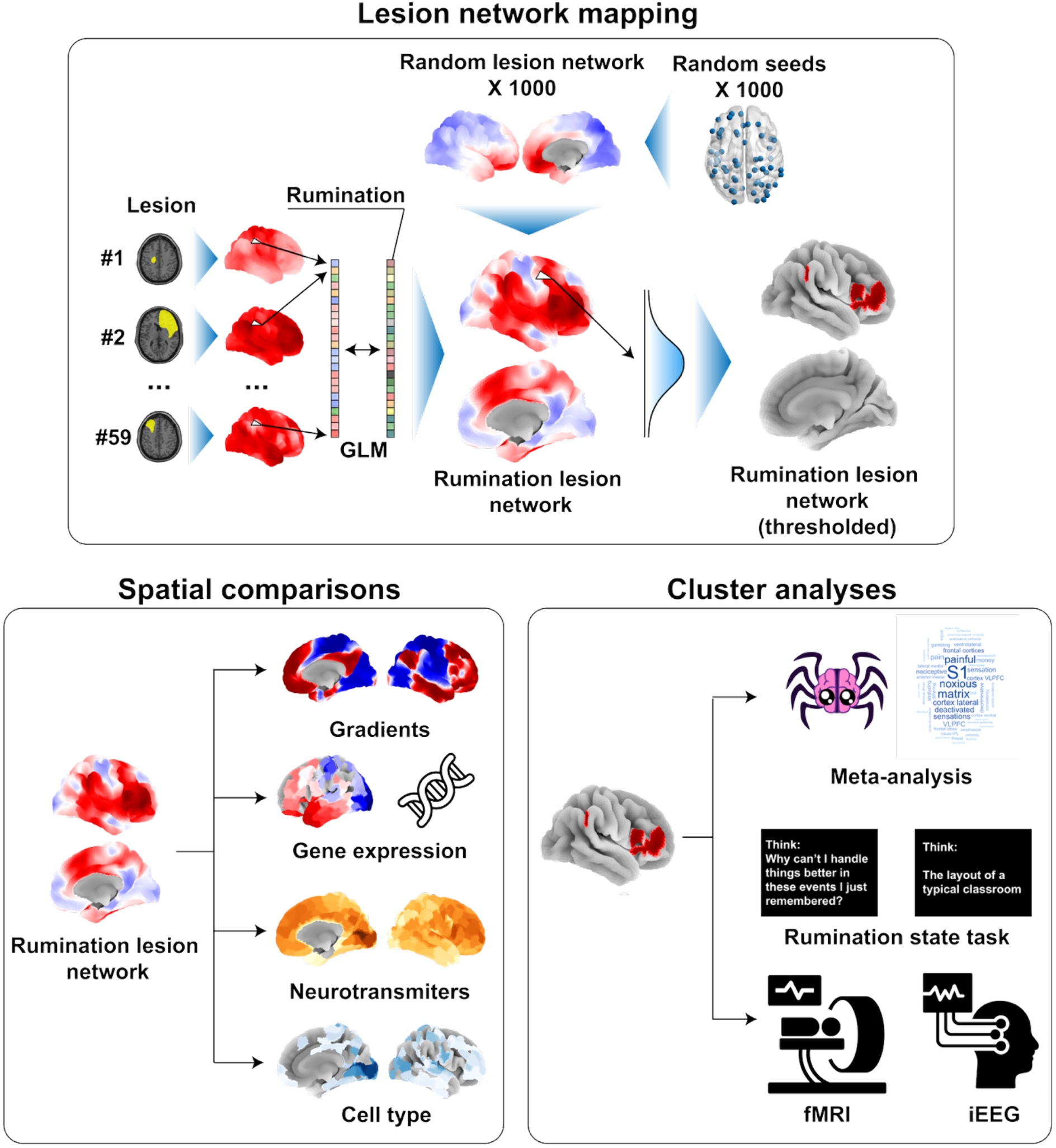
Flowchart of the lesion network mapping framework and multimodal characterization of the rumination-related brain circuit. Top: Lesion locations associated with depressive rumination were mapped onto whole-brain functional connectivity networks. A permutation-based approach, incorporating 1,000 random lesion networks and 1,000 random seeds, was used to construct null distributions and identify a rumination-related lesion network, which was subsequently thresholded to delineate significant regions. Bottom left: The identified rumination-related network was characterized through spatial comparisons with macroscale functional gradients, cortical gene expression profiles, neurotransmitter receptor distributions, and cell type maps. Bottom right: The resulting network was further investigated through meta-analytic functional decoding and experimental validation using a rumination-state task, incorporating functional magnetic resonance imaging (fMRI) and intracranial electroencephalography (iEEG). Together, these complementary analyses aimed to establish the spatial, molecular, and functional characteristics of the brain circuit associated with depressive rumination.

**Figure 2.**
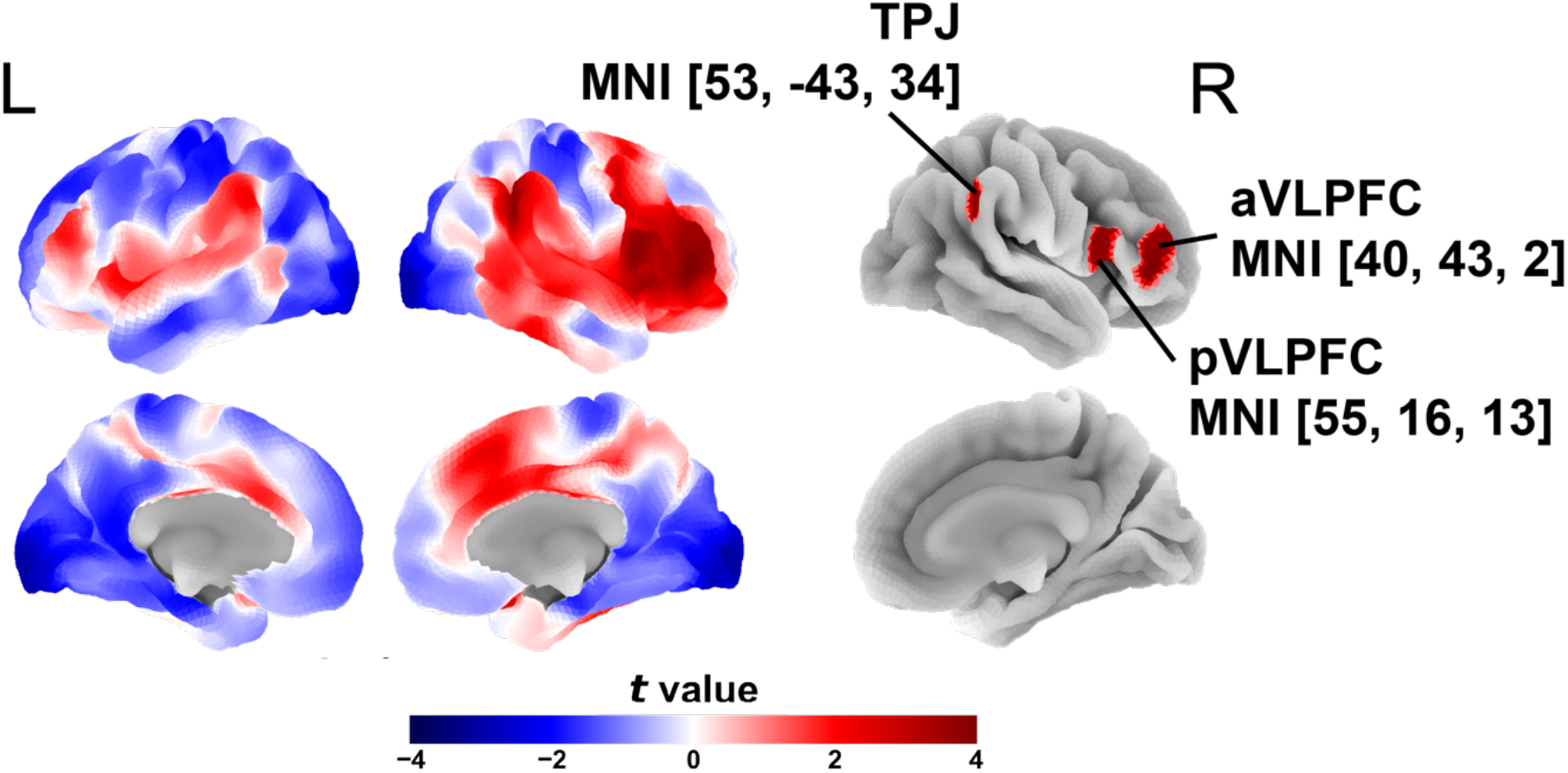
Spatial distribution of the rumination-related lesion network and its significant clusters. Left: Cortical surface representations of the rumination-related lesion network derived from lesion network mapping, displayed in lateral and medial views of both hemispheres. Right: Significant clusters identified using permutation-based statistical inference. Abbreviations: TPJ, temporal parietal junction; aVLPFC, anterior ventral lateral prefrontal cortex; pVLPFC, posterior ventral lateral prefrontal cortex.

**Table 1.** Included datasets.

| Dataset identifier | Setting | *n* | Diagnosis | Age (years) |  | Sex (female) |  |
|---|---|---|---|---|---|---|---|
| GLI-RUM | Cohort | 59 | Glioma | 44.07 (12.57) |  | 45.76% |  |
| DIRECT | Multi-center consortium | 3001 | MDD | MDD | HC | MDD | HC |
|  |  |  |  | 32.92 (13.15) | 32.35 (12.95) | 63.79% | 57.19% |
| RST-Suzhou | Cohort | 120 | MDD | 25.86 (7.60) | 28.95 (10.12) | 73.81% | 76.74% |
| RST-Beijing | Cohort | 41 | HC | 22.75 (3.68) |  | 52.5% |  |
| RST-iEEG | Cohort | 32 | Epilepsy | 27.38 (8.88) |  | 50% |  |

Age is reported as mean (SD).

### Relation with large-scale networks and gradients

We next sought to explore the relationship between the delineated rumination circuits and the brain network and the gradient structures ^31,38,39^. Because positive and negative values of the rumination circuit may reflect opposing functional roles in rumination, we analyzed the two components separately for their overlap with large-scale brain networks. We found that the negative rumination circuits mostly overlapped with the default mode network (DMN, *D* = 0.332) and visual network (VN, *D* = 0.380), while the positive circuit mostly overlapped with the ventral attention network (VAN, *D* = 0.365) and fronto-parietal network (FPN, *D* = 0.283)(Figure 3a). These results suggest that the positive and negative circuits may support distinct brain functions underpinning rumination. We found that the rumination circuit was significantly correlated with Gradient 3 (MR, Spearman’s rho = 0.507, *P_adjusted_* < 0.001)(Figure 3b). Further analyses revealed a clear organization along Gradient 3. Circuit values increased progressively along Gradient 3, transitioning from predominantly negative values at the lower end to positive values at the higher end.

**Figure 3.**
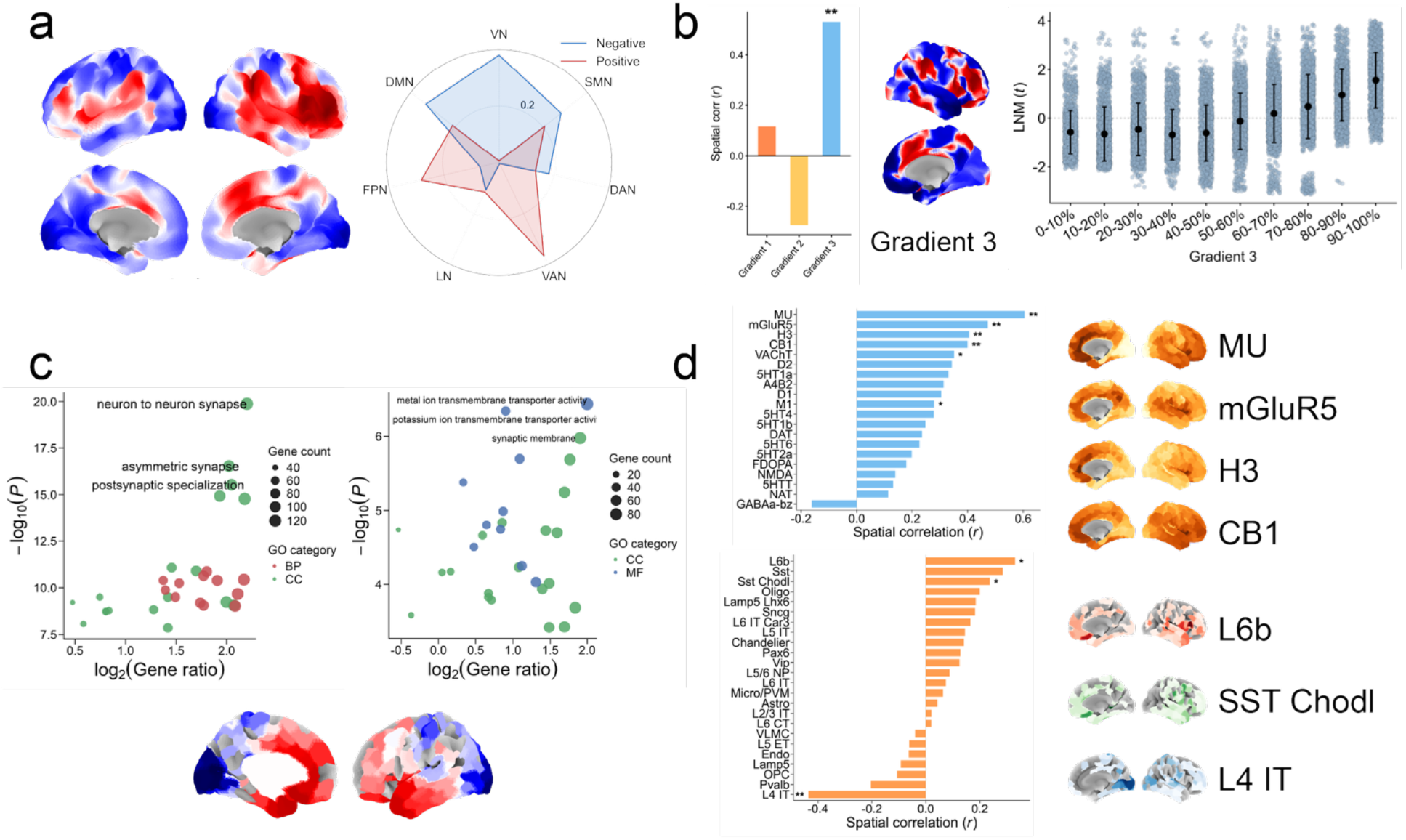
Multiscale spatial and molecular characterization of the rumination-related lesion network. (a) The rumination-related lesion network and its spatial overlap with the seven canonical functional networks. The radar plot illustrates the relative contributions of positive and negative network regions to each functional network. (b) Spatial correlations between the rumination-related lesion network and the principal functional gradients. Gradient 3 (the modulation–representation [MR] axis) showed the strongest spatial association with the rumination-related network. The scatterplot illustrates the distribution of LNM values across deciles of Gradient 3, revealing a progressive increase in LNM values along the gradient. (c) PLS and GO enrichment analyses of genes associated with the spatial distribution of the rumination-related network. Enriched BP, CC, and MF predominantly involved synaptic organization, neuronal communication, and ion transport. Dot size represents gene count, and statistical significance is expressed as −log₁₀(*P*). Cortical maps illustrate the spatial distribution of gene expression patterns associated with the network. (d) Spatial correlations between the rumination-related network and neurotransmitter receptor/transporter distributions (top), as well as cell-type profiles (bottom). Representative cortical maps illustrate the spatial distributions of selected neurotransmitter receptors and cellular markers. \*\**P* < 0.05, corrected for multiple comparisons; \**P* < 0.05, uncorrected. Abbreviations: LNM, lesion network mapping; GO, Gene Ontology; BP, biological processes; CC, cellular components; MF, molecular functions.

### The rumination circuit is associated with a certain gene expression profile, neurotransmitter, and cell type maps

To relate the cortical gene expression to parcel-wise rumination circuit values, we fitted partial least squares (PLS) models. PLS models were fitted on adult human cortical gene expression (AHBA ^40^, abagen ^41^, Schaefer-400 parcellation ^38^) and oriented so that the correlation between the first PLS parcel score and the mean rumination circuit value was non-negative. The resulting component maps define smooth molecular axes across the cortex for the rumination circuit.

The first component of the PLS model explained 37.74% of the variance. PLS1 gene expression weights were positively correlated with rumination circuit values (Spearman’s rho = 0.536, *P* < 0.001) (Figure S3). Gene ontology (GO) analysis highlighted synaptic organization and vesicle-related processes in the positive PLS1 gene set, and ion transport and channel activity in the negative set, with synaptic components enriched in both sets. We next investigated the association between the rumination circuit by relating the rumination circuit values and the established neurotransmitter profiles ^34^. We found significant positive associations between the rumination circuit profile and the μ-opioid receptor (MU; Spearman’s rho = 0.605, *P*_adjusted_ < 0.001), metabotropic glutamate receptor 5 (mGluR5; Spearman’s rho = 0.492, *P*_adjusted_ = 0.001), histamine H3 receptor (Spearman’s rho = 0.450, *P*_adjusted_ = 0.012), and cannabinoid receptor type 1 (CB1; Spearman’s rho = 0.418, *P*_adjusted_ = 0.018). Next, we explored the relation between the rumination circuit profile and the cell type maps ^35^. The rumination circuit showed a significant negative association with L4 IT, an excitatory neuron subtype (Spearman’s rho = −0.435, *P*_adjusted_ = 0.024). Positive associations were also observed for two other neuronal subtypes (L6b and Sst) before correction for multiple comparisons (L6b, Spearman’s rho = 0.330, *P*_unadjusted_ = 0.006; Sst, Spearman’s rho = 0.237, *P*_unadjusted_ = 0.034).

### The psychological processes related to the significant rumination circuit nodes

We generated the vertex-wise null distribution using randomly located coordinates. The *Z* scores were calculated according to the mean and standard deviation of the null distribution. We adopted a threshold of |*Z*| ≥ 3.29, revealing three significant rumination circuit nodes: anterior ventral lateral prefrontal cortex (aVLPFC), posterior VLPFC (pVLPFC), and temporal parietal junction (TPJ) (Figure 2). We next explored the psychological processes that are related to these nodes using Neurosynth. We found that these 3 clusters bear different psychological processes. aVLPFC node was associated with the sensation of pain and reward. pVLPFC node was associated with interoceptive processes, emotional information, and inhibition. TPJ node was response inhibition, consciousness, and navigation (Figure 4a).

**Figure 4.**
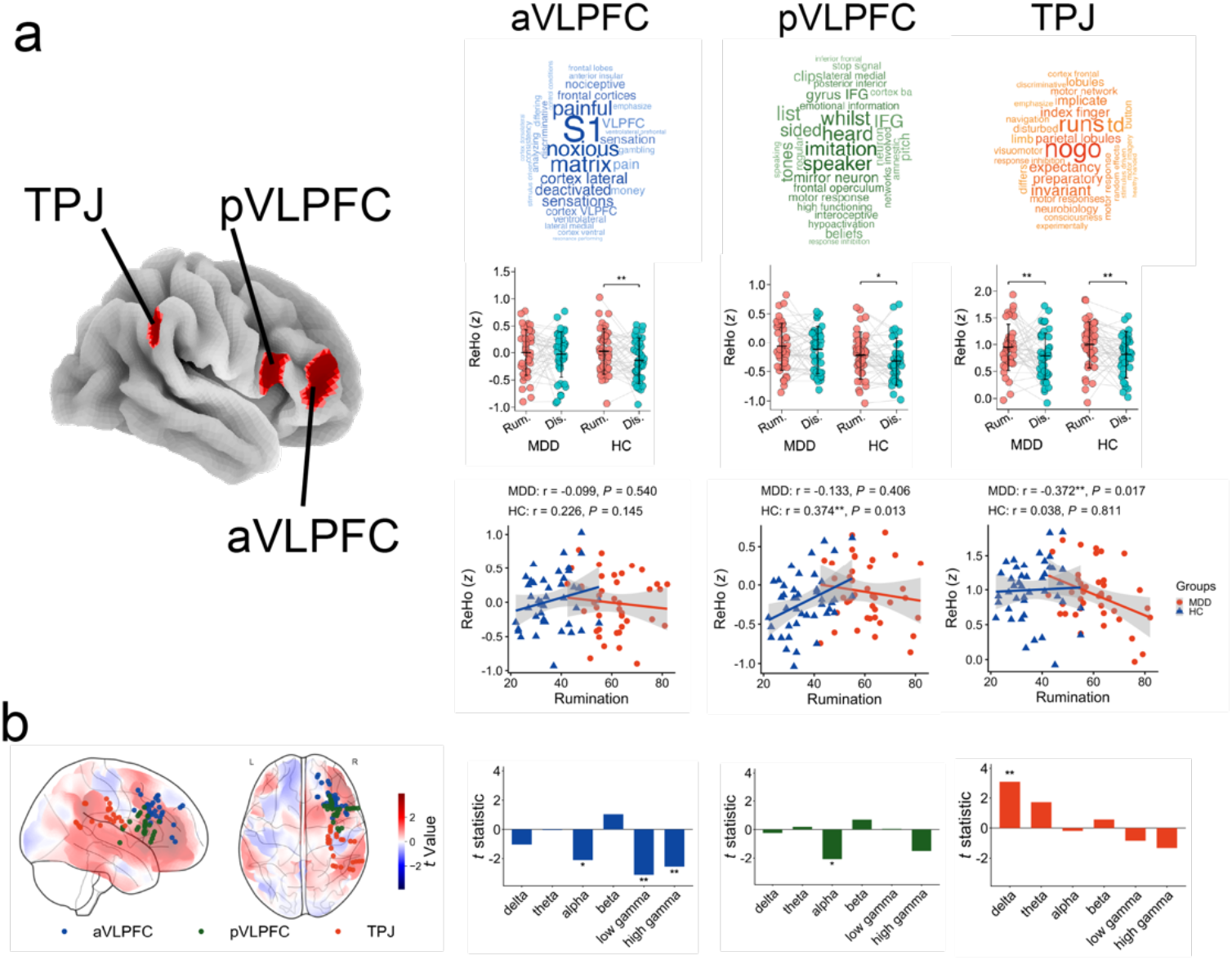
Functional characterization and cognitive relevance of brain regions associated with depressive rumination. (a) Functional decoding of the identified regions using Neurosynth, showing the most strongly associated cognitive and psychological terms. Regional differences in ReHo between patients with MDD and HC during rumination and distraction conditions. Associations between ReHo and rumination severity within each group. Each column represents a distinct region. (b) iEEG contacts included in the analyses. Highlighted in different colors for each brain region. Contact locations are overlaid on LNM maps. Frequency-specific differences in iEEG spectral power between rumination and distraction conditions across contacts located in different brain regions. Bar plots show *t* statistics comparing spectral power between conditions across six frequency bands. MDD, major depressive disorder; HC, healthy controls; Rum., rumination; Dis., distraction.

We next used an established rumination state task (RST) to validate the relevance of these critical nodes to rumination. In RST, participants were induced into a continuous rumination state and a control state (distraction). By contrasting rumination and distraction states, neural processes that are relevant to rumination could be revealed. We compared a widely used local fMRI metric, ReHo, a proxy for correlations among brain activity within a local brain region, between the rumination and distraction states. We found that the TPJ node showed significantly enhanced ReHo in the rumination state as compared to the distraction state in both individuals with MDD and HCs (MDD, *t*(41) = 2.826, *P*_adjusted_ = 0.022, Cohen’s *d* = 0.436; HC, *t*(42) = 3.258, *P*_adjusted_ = 0.006, Cohen’s *d* = 0.497). This effect was replicable in the RST-Beijing cohort, though results varied across scanners with marginal or non-significant findings while retaining consistent effect directions (IPCAS, *t*(39) = 1.884, *P*_adjusted_ = 0.201, Cohen’s *d* = 0.298; PKUGE, *t*(39) = 2.462, *P*_adjusted_ = 0.055, Cohen’s *d* = 0.389; PKUSIEMENS, *t*(39) = 3.016, *P*_adjusted_ = 0.013, Cohen’s *d* = 0.477). The TPJ’s ReHo showed a significant negative correlation with ruminative tendency in individuals with MDD (*r* = −0.372, *P* = 0.017), whereas no such effect was observed in HCs (*r* = 0.038, *P* = 0.811). Both aVLPFC (MDD, *t*(41) = 0.525, *P*_adjusted_ = 1.808, Cohen’s *d* = 0.081; HC, *t*(42) = 3.002, *P*_adjusted_ = 0.013, Cohen’s *d* = 0.458) and pVLPFC (MDD, *t*(41) = 1.246, *P*_adjusted_ = 0.638, Cohen’s *d* = 0.192; HC, *t*(42) = 2.310, *P*_adjusted_ = 0.078, Cohen’s *d* = 0.352) nodes showed enhanced ReHo in HCs while individuals with MDD showed insignificant but same direction effects. The enhancement of aVLPFC in HCs could be replicated in RST-Beijing cohort (IPCAS, *t*(39) = 3.244, *P*_adjusted_ = 0.002, Cohen’s *d* = 0.513; PKUGE, *t*(39) = 2.601, *P*_adjusted_ = 0.039, Cohen’s *d* = 0.411; PKUSIEMENS, *t*(39) = 3.244, *P*_adjusted_ = 0.002, Cohen’s *d* = 0.513), while the effect in pVLPFC could not be reproduced (IPCAS, *t*(39) = −0.126, *P*_adjusted_ = 2.702, Cohen’s *d* = − 0.020; PKUGE, *t*(39) = −1.918, *P*_adjusted_ = 0.187, Cohen’s *d* = −0.303; PKUSIEMENS, *t*(39) = 0.226, *P*_adjusted_ = 0.822, Cohen’s *d* = 0.036). We observed no significant association between ReHo and ruminative tendency in aVLPFC or pVLPFC, except for pVLPFC ReHo in HCs (*r* = 0.374, *P* = 0.013).

To further test whether the revealed critical rumination circuit nodes correspond to specific neural underpinnings of active ruminative processes, we examined iEEG data during an active ruminative state from a group of individuals with refractory epilepsy. We compared the mean oscillation band power of electrodes located in the same FreeSurfer segmentation of the rumination circuit nodes between the rumination and distraction states. We found significantly reduced low gamma (*t*(30) = −3.141, *P*_adjusted_ = 0.023, Cohen’s *d* = −1.147) and high gamma (*t*(30) = −2.85, *P*_adjusted_ = 0.045, Cohen’s *d* = −0.944) band power during rumination compared to distraction in the aVLPFC. We also found enhanced delta band power in the TPJ region during rumination as compared to the distraction states (*t*(32) = 3.106, *P*_adjusted_ = 0.024, Cohen’s *d* = 1.098)(Figure 4b). No significant band power alterations were found in the pVLPFC region. Thus, both the fMRI and iEEG data indicated that the aVLPFC and TPJ nodes of the rumination circuit may correspond to specific local neural activities.

### Consistent rumination circuit nodes FC profiles across groups and datasets

To characterize the network-level functional connectivity profiles of the aVLPFC, pVLPFC, and TPJ and assess their consistency across samples, we examined seed-based connectivity in participants with MDD, HCs from the RST-Suzhou cohort, and the RST-Beijing cohort. We found reproducibly enhanced TPJ-FPN FC during rumination as compared to distraction (RST-Suzhou: MDD, *t*(40) = 2.096, *P*_adjusted_ = 0.149, Cohen’s *d* = 0.348, HC, *t*(41) = 4.862, *P*_adjusted_ < 0.001, Cohen’s *d* = 1.011; RST-Beijing: IPCAS, *t*(39) = 5.249, *P*_adjusted_ < 0.001, Cohen’s *d* = 0.840, PKUGE, *t*(39) = 3.735, *P*_adjusted_ = 0.004, Cohen’s *d* = 0.583, PKUSIEMENS, *t*(39) = 3.639, *P*_adjusted_ = 0.006, Cohen’s *d* = 0.638)(Figure 5 and S6). We repeatedly observed enhanced FCs between TPJ and right VLPFC, inferior parietal lobule (IPL), and left medial temporal lobe (MTL). We also found reproducible reduced FCs between TPJ and the left dorsal medial prefrontal cortex (DMPFC) and VLPFC. We also found a reproducible enhanced aVLPFC-FPN FC during the rumination state (RST-Suzhou: MDD, *t*(40) = 1.758, *P*_adjusted_ = 0.302, Cohen’s *d* = 0.300, HC, *t*(41) = 3.384, *P*_adjusted_ = 0.007, Cohen’s *d* = 0.619; RST-Beijing: IPCAS, *t*(39) = 3.531, *P*_adjusted_ = 0.008, Cohen’s *d* = 0.562, PKUGE, *t*(39) = 3.212, *P*_adjusted_ = 0.019, Cohen’s *d* = 0.428, PKUSIEMENS, *t*(39) = 1.841, *P*_adjusted_ = 0.515, Cohen’s *d* = 0.282). Reproducible reduced FCs between aVLPFC and the left DMPFC and enhanced FCs between aVLPFC and the dorsal precuneus and the right retrosplenial cortex (RSC), although these effects did not reach significance in some cases. We observed no reproducible effect in the pVLPFC. These findings suggest that aVLPFC and TPJ may relate to rumination through an enhanced coupling with FPN.

**Figure 5.**
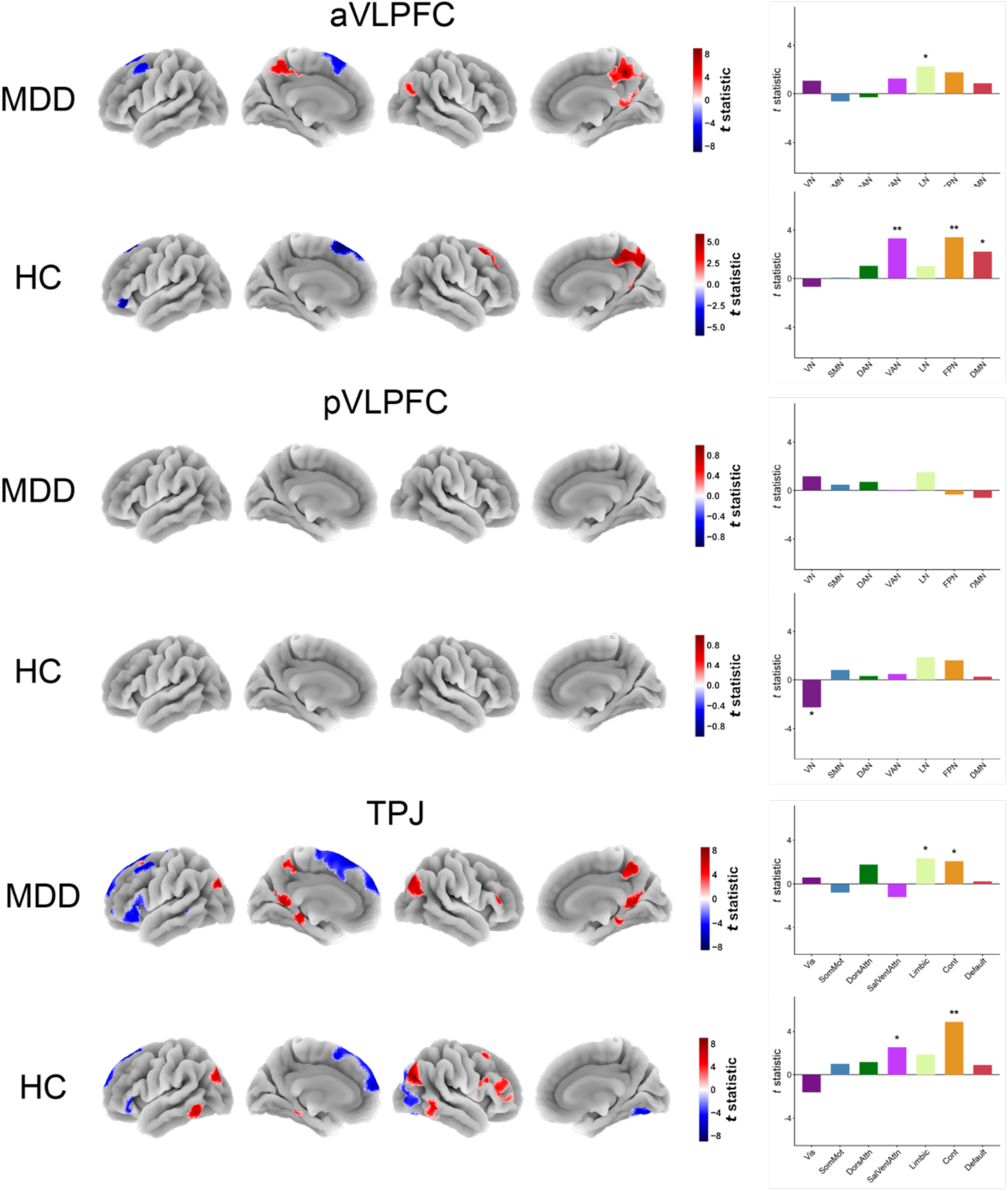
Whole-brain FC patterns associated during the rumination condition compared with the distraction state. Bar plots denote the network-wised differences. Abbreviations: TPJ, temporal parietal junction; aVLPFC, anterior ventral lateral prefrontal cortex; pVLPFC, posterior ventral lateral prefrontal cortex; MDD, major depressive disorder; HC, healthy controls. VN, visual network; SMN, somato-motor network; DAN, dorsal attention network; VAN, ventral attention network; LN, limbic network; FPN, fronto-parietal network; DMN, default mode network.

## Discussion

In this work, we integrated lesion network mapping with functional MRI and iEEG across five independent datasets to identify a distributed brain circuit associated with rumination. The circuit was highly consistent across normative connectomes derived from individuals with MDD and healthy controls and remained robust to GSR. Its positive and negative components showed distinct relationships with large-scale functional networks. Its spatial profile was associated with cortical functional gradients, gene expression, and regional molecular and cellular maps. Empirical null-model testing identified three significant nodes in the anterior and posterior VLPFC and TPJ. Across three independent datasets, we showed that these nodes exhibited specific changes in ReHo, intracranial oscillatory power, and FC with FPN during an active ruminative state. Together, these findings provide a lesion-informed account of the distributed neural architecture of rumination and nominate the aVLPFC and TPJ as candidate targets for future neuromodulation studies.

Here, we extend previous neuroimaging findings ^12,13,42–45^ implicating an association between rumination and complex functional coupling among DMN, FPN, and attention networks by leveraging LNM to provide causal evidence. We found that the negative component of the lesion-derived map overlapped primarily with the DMN and visual network, whereas the positive component overlapped predominantly with the VAN and FPN. One possible interpretation is that the negative component supports rumination, as lesions connected to these regions were associated with lower ruminative tendency, whereas the positive component constrains rumination, as lesions connected to these regions were associated with higher ruminative tendency. Existing studies using RIT found that DMN regions were activated, whereas FPN, VAN, and DAN regions were deactivated when participants were induced into rumination ^7,9,10^. Previous cognitive frameworks of rumination have implicated that rumination may reflect difficulty inhibiting or updating components in working memory ^46–49^. Together, these findings suggest that rumination may arise from an imbalance between networks that support internally generated thought and those that enable its flexible regulation.

The spatial associations with functional gradients and molecular and cellular maps further contextualize the rumination circuit within the cortex’s multiscale organization. Functional coupling among motifs on the human brain surface is reliably organized along three dominant gradients which together provide a low-dimensional framework for describing large-scale cortical architecture ^50^. The three established prominent gradients are the sensory-association (SA) axis, the visual-somatosensory axis (VS axis), and the modulation–representation (MR) axis ^51^. The SA axis is anchored at one end by primary unimodal cortex (VN and SMN) and at the other end by high-order association cortex (i.e., DMN) ^31^, explaining the most variance in FC. The VS axis spans between the VN and SMN, capturing a continuous transition in cortical specialization for visual versus somatomotor processing. The MR axis distinguishes cortical regions primarily involved in modulation, including frontoparietal control and attention networks, from those supporting representation, including default, sensory, and visual networks ^52,53^. Here, we found a significant positive spatial correlation between the rumination circuit and the MR axis. The MR gradient captures an information-processing hierarchy that is critical to complex cognition, with higher-order association and control regions at its apex. Supporting this, we recently found that the active rumination state is characterized by reduced gradient values within the DMN regions ^32^. Turnbull et al. ^54^ found that the spatial pattern of off-task thought aligned selectively with the MR axis, but not with the remaining two gradients, supporting the relevance of this axis to spontaneous thought. Together, these findings suggest that the rumination circuit is situated along a cortical axis relevant to the modulation and representation of thought.

To bridge gaps across levels of biological systems, we related adult cortical gene expression (AHBA, abagen) to the rumination LNM profile using a supervised PLS approach. Genes with positive PLS1 weights were enriched for synaptic organization and vesicle-related processes, whereas those with negative weights were enriched for ion transport and channel activity, with synaptic components represented in both sets. We further found that the rumination circuit values were positively associated with the distributions of μ-opioid, mGluR5, histamine H3, and CB1 receptors, and negatively associated with the L4 IT excitatory neuron map. Together, these observations characterize the rumination circuit across molecular, neurochemical, and cellular scales and motivate hypotheses concerning how synaptic signaling and cellular architecture may contribute to its functional organization. The opioid signaling could be influenced by childhood trauma and lead to dysfunctional response styles, including depressive rumination ^55^. Metabotropic glutamate receptors play a critical role in mice’s cognitive flexibility ^56–58^, which is closely related to humans’ difficulty disengaging from negative thoughts in ruminative processes ^59,60^. The histaminergic molecular architecture is involved in emotion regulation, salience processing, and cognitive control ^61^. The CB1 receptor deletion in mice increases anxiety-like behavior and disrupts self-care and exploratory behavior ^62^. L4 IT cells are mainly distributed in layer 4, which is highly specialized for processing incoming sensory information ^63^. The negative spatial association between L4 IT cell-type abundance and rumination circuit values may reflect a distinction between the sensory-input and feedforward processing functions of L4 IT neurons and the higher-order associative and self-referential processes implicated in rumination ^1,7^. Our findings suggest that the spatiomolecular axes, alongside their molecular and cellular maps, may provide a scaffold within which the rumination circuit is embedded ^51^.

The correlation between significant rumination nodes and meta-analytic functional neuroimaging maps from Neurosynth indicated that brain regions significantly related to rumination were strongly associated with reward, introspection, and inhibition processes. Independent fMRI cohorts showed that these nodes’ local BOLD activity metrics (ReHo) were enhanced during an active ruminative state. TPJ’s ReHo during rumination was negatively related to the ruminative tendency in individuals with MDD. iEEG evidence showed that aVLPFC’s gamma-band oscillation power was reduced and TPJ’s delta-band power was enhanced during rumination. Further whole-surface FC analyses showed increased FC between both aVLPFC and TPJ and the FPN. Replicable clusters included the left DMPFC, dorsal precuneus, and right RSC. These results suggest that the VLPFC and TPJ may play critical roles in the rumination process through both altered local neural activity and functional coupling to the FPN. The VLPFC may be distinguished by a dense, interdigitated arrangement of association networks that facilitates cross-network integration in support of abstract and complex cognition ^64^, while the TPJ is a supramodal association region that is involved in both attentional and social processes ^65,66^. It should be noted that VLPFC and TPJ are two prominent nodes in a framework of inhibitory control over thought ^67^. Within this framework, VLPFC involvement may reflect attempts to regulate ruminative content, whereas TPJ involvement may relate to attentional capture by internally generated thoughts and the subsequent recruitment of control. Their increased coupling with the FPN may therefore reflect coordination between internal attentional processing and regulatory demands during rumination. Supporting this, we found a significant negative association between ReHo in TPJ and the ruminative tendency in individuals with MDD. This interpretation remains tentative, however, because our task induced rumination rather than explicitly testing thought suppression, and increased functional connectivity does not establish either the direction or success of inhibitory control.

Our iEEG findings provide rare evidence for the neural substrate of rumination. Previous scalp EEG studies have found altered oscillation power and coherence during an active ruminative state ^68,69^. Our prior iEEG study showed that the delta/theta band power was enhanced in the precuneus region ^18^. Our results further showed that iEEG could uncover nuanced differences in the local neural underpinnings of distinct brain nodes of the rumination circuit. Although the VLPFC and TPJ showed similar patterns in the fMRI metrics (ReHo and FC), their electrophysiological features are different (enhanced delta band power in TPJ and reduced gamma band power in VLPFC).

A major strength of our study is replicating the fMRI results across sites (Rum-Suzhou) and repeated assessments (Rum-Beijing), providing complementary evidence for their generalizability and temporal robustness. This is particularly important given persistent reproducibility challenges in psychiatric neuroimaging ^70^. Cao et al. recently reported low cross-site consistency of morphometric differences across five psychiatric disorders, despite harmonized processing and extensive sensitivity analyses ^71^. Against this background, our findings converge across sites, reducing concerns that they depend on a particular recruitment setting or local acquisition conditions, while their recurrence across repeated assessments supports stability beyond a single measurement occasion. Together, these complementary forms of replication strengthen confidence in the robustness of the reported effects and provide a firmer foundation for subsequent mechanistic investigation and validation.

The brain regions identified here could serve as potential targets for future neuromodulation treatments focused on rumination, a viable treatment target that could be especially effective for certain groups of patients ^72^. Despite existing psychotherapeutic avenues ^73^, neuromodulation remains a promising therapeutic approach. Previous studies have provided preliminary evidence that neuromodulation can alleviate rumination and related negative thought processes, although substantial scope for improvement remains. Cathodal tDCS targeting bilateral inferior parietal nodes of the default mode network reduced negative past-oriented mind-wandering without altering overall mind-wandering frequency ^74^. In a retrospective study of treatment-resistant depression, rTMS was associated with improvements in both rumination and depressive symptoms, but the reduction in rumination was smaller than that in depression ^75^. These findings support the potential of neuromodulation while highlighting the need to optimize interventions specifically for rumination. One possibility is that existing stimulation targets do not optimally engage the neural circuits underlying rumination. The candidate targets identified using our approach therefore offer a promising direction for future research, with prospective, sham-controlled trials needed to determine whether stimulating these sites produces greater and more durable reductions in rumination.

Several limitations should be considered. First, the lesion-derived circuit was identified in a cohort of patients with diffuse glioma, with uneven anatomical coverage. Tumor-related effects and potential functional reorganization may influence the observed associations ^76^, limiting generalizability to other lesion etiologies and individuals without structural brain lesions. Second, the random-coordinate null model also does not fully reproduce the spatial extent and anatomical distribution of glioma lesions, leaving scope for further validation using complementary null models ^37^. Finally, neither lesion connectivity nor task-related activity establishes whether stimulation of a candidate node would reduce rumination or identifies the appropriate stimulation protocol. Prospective, sham-controlled trials with rumination-specific outcomes are therefore needed to establish the therapeutic relevance of the aVLPFC and TPJ.

Overall, this study integrated lesion network mapping, functional MRI, and intracranial electrophysiology to delineate a distributed brain circuit associated with rumination. The circuit contained peaks in the right VLPFC and TPJ, which could serve as potential novel brain stimulation targets for rumination. The circuit aligned with the cortical modulation–representation gradient and showed spatial associations with molecular and cellular profiles. Complementary evidence from independent datasets supported the functional relevance of the aVLPFC and TPJ during active rumination, implicating distinct local neural dynamics alongside increased coupling with the frontoparietal network. These findings provide a lesion-informed framework for understanding the neural organization of rumination and identify candidate targets for future neuromodulation research.

## Methods

### Overview

This study consisted of five independent datasets (GLI-RUM, DIRECT, RST-Suzhou, RST-Beijing, RST-iEEG; total *n* = 3242). DIRECT was a publicly available, multi-site consortium (https://rfmri.org/DIRECTPhase2) ^77^. The remaining datasets were locally collected, and RST-Suzhou ^15^ and RST-Beijing ^14^ have been made publicly available (RST-Suzhou: https://osf.io/wts7f/overview; RST-Beijing: https://doi.org/10.57760/sciencedb.o00115.00002). We used GLI-RUM and DIRECT to map lesions to a common brain network and delineate its correlation with rumination. RST-Suzhou, RST-Beijing, and RST-iEEG ^18^ were used to validate the functional relevance of the revealed rumination circuit to an induced active ruminative state. In each dataset, participants provided informed consent, and the institutional review board approved the data collection.

GLI-RUM was collected at Huashan Hospital, Shanghai, China. We analyzed data from 59 participants diagnosed with diffuse glioma. All participants completed the ruminative response scale (RRS) ^11^ and Beck depression inventory (BDI) ^78^. We manually segmented lesions based on structural MRI images, spatially normalized to the Montreal Neurological Institute (MNI) 152 atlas space, and binarized them.

DIRECT consists of harmonized resting-state functional MRI (rs-fMRI) data of individuals with major depressive disorder (MDD; *n* = 1660) and healthy controls (HC; *n* = 1341). Details can be found elsewhere ^77^. Briefly, researchers aggregated 23 case-control-designed rs-fMRI datasets and adopted a coordinated preprocessing pipeline (DPABISurf ^79^) to alleviate the batch effect.

RST-Suzhou was collected at Guangji Hospital in Suzhou, Jiangsu, China, from December 2021 to September 2022. In this dataset, individuals with MDD and HCs completed a validated rumination state task (RST) ^14,15^, in which they were induced into a continuous, active ruminative state and a control state (distraction). RST has been found to be able to reliably characterize the functional network underpinnings of depressive rumination ^14,16,17^. This study analyzed only the rumination and distraction states. Details regarding RST are provided in the supplementary information. Participants’ ruminative tendency was measured using RRS.

RST-Beijing was collected from the local community near the Institute of Psychology, Chinese Academy of Sciences (IPCAS). A group of healthy adults completed the RST on 3 different scanners at two sites (one GE MR750 scanner at IPCAS, one GE MR750 and one SIEMENS Prisma scanner at Peking University).

RST-iEEG was collected at Huashan Hospital, Shanghai, China. A group of individuals with refractory epilepsy (*n* = 21) who had stereo-electroencephalography implants for presurgical evaluation was recruited and completed RST during intracranial epilepsy monitoring.

### Lesion network mapping

Lesion network mapping was performed using the GLI-RUM and DIRECT datasets to identify brain networks causally associated with individual differences in rumination. This network was identified in accordance with previously published methods using custom MATLAB and Python codes ^21,23,80^. Two normative connectomes of individuals with MDD (*n* = 1574) and HCs (*n* = 1308) from DIRECT who survived the quality control (QC) were used to generate rs-fMRI functional connectivity (FC) maps for each glioma lesion of the GLI-RUM dataset (see the supplementary information for the QC procedure). Rs-fMRI data were preprocessed in accordance with prior work ^77^, and details were provided in the supplementary information. The site effect was controlled using comBat. Whole-surface FC was computed for each glioma lesion as illustrated in Figure 1. For each lesion, we extracted the mean BOLD time series within the lesion mask and then computed whole-surface FC maps by correlating the lesion mean BOLD time series with all vertices across the brain surface. We then averaged the FC maps across all participants in the cohort (individuals with MDD or HCs in DIRECT) to generate a lesion-specific normative FC map. Once all the normative FC maps were generated, these FC maps were correlated with the corresponding individual RRS scores using a general linear model (GLM), with age, sex, education level controlled. This analysis yielded a circuit map representing the connectivity of lesions that modify rumination.

Next, this circuit was subject to statistical testing (Figure 1). This procedure mimicked that described above, except that FC maps were derived from spheres (5 mm radius) that were located in randomly generated coordinates. The number of contributing coordinates was held identical to the number of gliomas (*n* = 59). We then fitted vertex-wise GLMs to calculate the null rumination-related circuits. We repeated this procedure 1,000 times to generate 1,000 null rumination-related circuits. We used the empirical null distribution to assess whether the observed circuit was associated with the psychological process of interest. For each condition, a *Z*-score map was generated by subtracting the mean of the 1,000 null samples from the observed FC value and dividing by its standard deviation. This standardization was performed independently at each vertex.

### Convergence and comparison with previous network models

We also tested the relationship between the rumination circuit delineated here and previous surface network models. We first explored the correspondence between the present rumination circuit and the well-established Yeo 7 network structure ^39^. Given that the positive and negative components of the rumination circuit may have distinct functional implications, potentially supporting and opposing rumination, respectively, we first separated the circuit into positive and negative maps. We then quantified the spatial overlap of each component with each of the seven canonical functional networks from the Yeo atlas using the Dice coefficient.

We next explored the spatial similarity between the rumination circuits and the principal gradient maps. The gradient maps were extracted from the neuromaps toolbox ^81^. We focused on three reproducible axes—sensory-association (SA), visual-somatosensory (VS), and modulation-representation (MR). We calculated vertex-wise spatial correlation using Spearman correlation and determined significance using the spin test (10000 iterations). The multiple comparisons for the 3 axes were adjusted using Bonferroni correction. The vertices were divided into ten approximately equal-sized bins according to gradient-value deciles, spanning the lowest (0–10%) to highest (90–100%) gradient values. Within each bin, we calculated the mean lesion network mapping value and its standard error, providing a descriptive profile of the rumination circuit along the gradient.

### Association between the rumination circuit and gene expression profiles

We used the AHBA ^40^ microarray data processed with abagen ^41^ and parcellated according to the Schaefer-400 atlas (7-network, 2 mm, MNI 152). The abagen AHBA processing pipeline was run following the published recommendations ^82^. Analyses were restricted to the 200 left-hemisphere parcels, with right-hemisphere tissue samples mirrored to the left hemisphere before regional assignment. We applied a customized sample-filtering procedure to exclude unsuitable tissue samples, including samples distant from the cortex. The workflow specified thresholds of 3 for donor coverage and 0.5 for gene stability. The resulting regional expression matrix was reindexed to match the left-hemisphere parcel labels, with missing parcels retained as missing values. The rumination circuit values were averaged within the corresponding parcels. Before partial least squares (PLS) analysis, we excluded parcels with missing imaging or gene expression values, and separately standardized each gene’s expression profile and the regional imaging vector across retained parcels.

We then fitted PLS regression using the standardized gene expression matrix *X* as predictors and the regional rumination circuit vector *y* as the response (R package *pls* ^83^; 10 components). Subsequent analyses focused on the first component (PLS1), yielding parcel scores *t* = *Xw* and gene weights *w*. The component was oriented so that corr(*t*, *y*) ≥ 0, such that positive scores indicated parcels whose multigene expression profiles aligned with higher rumination circuit values. Gene–circuit coupling was summarized by the in-sample Pearson correlation between *t* and *y* and its square.

To assess the stability of gene contributions to this component, we resampled parcels with replacement 1,000 times, preserving the pairing between expression and imaging values, and refitted a single-component PLS model to each resample. Bootstrap weight vectors were sign-aligned with the original weight vector. For each gene, we estimated the standard error as the standard deviation of its bootstrap weights and calculated a bootstrap ratio by dividing its original weight by this standard error. Genes with bootstrap ratios > 3 and < −3 formed the positive and negative gene sets, respectively.

To characterize the biological processes and functions associated with these gene sets, we performed GO over-representation analysis separately for each set using *clusterProfiler* ^84^. Analyses covered biological process, molecular function, and cellular component terms, using the default background of genes represented in the corresponding GO annotation database. *P* values were adjusted using the Benjamini–Hochberg procedure separately within each gene set and ontology. Thresholds for unadjusted *P* values, adjusted *P* values, and *q* values were each set to 0.05.

### Spatial correlation with the neurotransmitter and cell type maps

We related the rumination circuit to 20 neurotransmitter maps ^34^ and 24 cell type expression maps ^35^ using the bilateral Schaefer-400 parcellation. Neurotransmitter and circuit values were averaged within parcels, and cell type maps were aligned by atlas parcel names. Spatial associations were quantified using Spearman correlation across parcels with complete data (400 for neurotransmitter maps; 339 for cell type maps). Statistical significance was evaluated against 5,000 bilateral spherical rotations of the circuit map with BrainSpace ^85^, using two-sided permutation tests. *P* values were corrected using Benjamini–Hochberg FDR at 0.05.

### Neurosynth functional decoding

To characterize the functional associations of the selected rumination-related cluster, we used Neurosynth ^86^ and NiMARE’s discrete Neurosynth decoder ^87^. The surface cluster mask was projected into MNI152 volumetric space, binarized and resampled to the Neurosynth analysis grid using nearest-neighbor interpolation. Studies reporting at least one activation coordinate within the mask were selected and compared with the remaining studies in the database. Term–activation associations were evaluated using the Neurosynth chi-square framework, with the default annotation threshold of 0.001 and term prior probability of 0.5. Terms were ranked by their reverse-inference posterior probability, reflecting the probability of a term given activation within the region.

### Rumination state task

Participants from the RST-Suzhou, RST-Beijing, RST-iEEG underwent a rumination state task, which has been validated in both healthy individuals ^14,16^ and patients with MDD ^17^. In general terms, this task consists of four sessions: a resting state, a negative autobiographical memory state, a rumination state, and a distraction state. The distraction state was included here as a control condition for two reasons. First, the distraction condition is different from the rumination condition on many dimensions. Second, it can prevent participants from entering spontaneous rumination. Each session lasted for 8 minutes. This study analyzed only the fMRI data from the rumination and distraction states. The order of these two sessions was counterbalanced across participants. During the rumination condition, participants were instructed to reflect on themselves according to prompts such as “Think: Why can’t I handle things better in these events I just remembered?” or “Think: Why do I encounter these events other people don’t?”. In contrast, they were instructed to imagine and focus on specific external topics using prompts such as “Think about: A train stopped at a station” or “Think about: The layout of a typical classroom” during the distraction state. All prompts were adapted from the rumination inducement task ^9,88^ and RRS ^89^. A total of 4 prompts lasting 2 minutes were displayed consecutively without inter-stimulus intervals in each state. For details regarding this task, please refer to our prior publications ^14,16,17^. Participants also reported their momentary emotional states at the end of each session. These manipulation-check data have been fully reported and validated in our prior publications using the same task ^17^. Briefly, previous findings showed significantly increased negative affect after the rumination condition, whereas emotional states returned to baseline after the distraction condition.

### Local fMRI metrics

We adopted a preprocessing pipeline that was identical to the pipeline used for the DIRECT dataset, with one exception: an additional covariate was included to control for the effect of the stimuli. Regional homogeneity (ReHo) ^90,91^ was picked to denote local neural activity for its superior reliability ^92^ and established neuropathological relevance ^93^. We calculated the so-called “2dReHo” using the following procedure. For each cortical vertex, Kendall’s coefficient of concordance (KCC) was computed across the BOLD time series of the vertex and its six immediate neighbors, defined by the surface mesh topology. The resulting KCC value was assigned to the central vertex as its ReHo value. This procedure was repeated across all vertices in both hemispheres to generate individual ReHo maps. For subsequent statistical analyses, we standardized each map by subtracting the mean ReHo across the entire cortical surface from each vertex’s value and dividing by the corresponding standard deviation. We extracted the mean ReHo values from each of the 3 significant clusters (anterior ventrolateral prefrontal cortex [aVLPFC], posterior ventrolateral prefrontal cortex [pVLPFC], and temporal parietal junction [TPJ]). We fitted a GLM to conduct paired *t*-tests on mean 2dReHo between the rumination and distraction states, controlling for head motion (mean FD_Jenkinson_). Multiple comparisons regarding the 3 significant clusters were controlled with Bonferroni correction. We further explored the association between ReHo and the individual ruminative tendency (RRS scores) using Pearson correlation.

### Comparing whole Surface FC maps between the rumination and distraction states

We extracted the mean preprocessed BOLD time series from the 3 significant clusters during the rumination and distraction states and then calculated vertex-wise Pearson correlations to create a whole-surface FC profile. Next, a vertex-wise GLM was fitted to perform paired *t*-tests, with head motion (mean FD_Jenkinson_) controlled. We adjusted for multiple comparisons with a permutation test (5000 iterations) using threshold-free cluster enhancement (TFCE) ^94^, which preserves reproducibility while controlling false-positive rates ^92^. We also extracted mean network FCs using the Schaefer-400 parcellation, grouped according to the Yeo 7 network atlas. Similarly, paired *t*-tests were performed controlling for head motion. Multiple comparisons were adjusted with FDR.

### iEEG preprocessing and spectral power analyses

FreeSurfer (v 6.0.0) ^95^ was used to align post-implantation CT brain scans from each patient, showing the location of the intracranial electrodes, to their pre-operative structural T1 MRI scans. Electrode positions were manually marked using FieldTrip (v 20251201) ^96^. We first determined the correspondence between the FreeSurfer segmentation and the significant clusters. We then assigned electrodes to an ROI according to the FreeSurfer segmentation, which we confirmed through independent expert visual inspection.

We first down-sampled the original iEEG signals to 1000 Hz and then band-pass filtered from 0.15 to 200 Hz using a two-pass Butterworth filter (ft_preprocessing.m in the FieldTrip toolbox ^96^). We notch-filtered the raw data at 50 Hz and its harmonics. We applied a bipolar montage to all depth electrodes by taking bipolar derivations of consecutive channels.

We visually inspected the raw iEEG data and their corresponding power spectrum to rule out channels containing signal artifacts or interictal epileptiform activity. We then applied an automatic removal procedure to the downsampled raw iEEG data. We obtained the iEEG data envelope using a Hilbert transform. The time points with epileptic discharges were identified according to the following criteria: (1) The envelope of the unfiltered signal exceeded the baseline by four standard deviations, where the baseline represents the signal’s overall mean value. (2) The filtered signal’s envelope, bandpass-filtered between 25 and 80 Hz, exceeded the baseline by five standard deviations, in line with prior research ^97^. We excluded channels with more than 50% of time points identified as noise from further analysis. The timing of all epileptiform activities and artifacts was stored for the following analysis.

We obtained a power spectrum using five-cycle Morlet wavelets at 76 logarithmically spaced frequencies from 0 to 150 Hz. This time-frequency transformation was performed using ft_freqanalysis.m in the FieldTrip toolbox. We excluded 1 second before and after the time points identified as noise in the automatic removal procedure described above. To control baseline inter-subject variability in the power spectrum, we then *Z*-scored the power values using the channel-level average power across all conditions of interest (e.g., rumination and distraction). We obtained these Z-scores by first subtracting the mean power and then dividing by the standard deviation (SD) across the conditions of interest. Finally, the mean powers of 5 frequency bands (delta, 1-4 Hz; theta, 4-8 Hz; alpha, 8-12 Hz; beta, 12-30 Hz; low gamma, 30-70 Hz; high gamma, 70-150Hz) were obtained by averaging all *Z*-standardized power value time series across all corresponding frequencies.

Condition-related differences in spectral power were assessed using linear mixed-effects models fitted separately for six frequency bands (delta, theta, alpha, beta, low gamma and high gamma). Each model included condition as a fixed effect and random intercepts for participants and electrodes nested within participants, accounting for multiple electrodes per participant and repeated measurements across conditions. Models were fitted using restricted maximum likelihood in MATLAB (*fitlme*). Distraction was the reference condition; thus, the condition coefficient represented the difference in power between rumination and distraction. Non-finite power values were excluded separately for each frequency band. We evaluated condition effects using two-sided tests of the fixed-effect coefficients, reporting estimates, standard errors, and 95% confidence intervals. *P* values were adjusted for multiple comparisons across the six frequency bands using the FDR procedure.

## Author Contributions

Concept and design: XC, ZF, C-GY. Acquisition of data: ZF, Y-YC, H-JL, XC, C-GY. Analysis or interpretation of data: XC, ZF, C-GY, T-FY. Drafting of the manuscript: XC, ZF. Critical revision of the manuscript for important intellectual content: XC, ZF, C-GY, T-FY, Y-YC, H-JL, T-CH, DY, X-YW, X-NZ, YZ, Z-WY, W-JZ. Statistical analysis: XC. Administrative, technical, or material support: ZF, XC. Supervision: T-FY, C-GY, XC.

## Data and code availability

The RST-Suzhou dataset is available through the OSF platform (https://doi.org/10.17605/OSF.IO/WTS7F). The RST-Beijing dataset is available through the Psychological Science Data Bank (https://doi.org/10.57760/sciencedb.o00115.00002). The DIRECT dataset is available through the Psychological Science Data Bank (https://doi.org/10.57760/sciencedb.psych.00689). The GLI-Rum dataset is not publicly available but may be shared with an institutional data use agreement and institutional review board approval. All codes related to the analysis and plotting of the present study have been openly shared (https://github.com/XiaoChenPhD/rumination_lesion_network_mapping/tree/main).

## Conflict of interest

The authors declare no conflicts of interest.

## Acknowledgements

This work was supported by the National Science & Technology Major Program on "Brain Science and Brain-Inspired Research" (No. 2026ZD0222303), the National Natural Science Foundation of China (No. 32300933).

## Additional Information

The authors used generative AI tools to help reorganize and copy-edit the manuscript. The tool was not used to generate scientific content, conduct or interpret analyses, draft results, or write the substantive sections of the manuscript. The authors reviewed and verified all AI-assisted text and take full responsibility for the integrity and accuracy of the content.

## Notes

### Competing Interest Statement

The authors have declared no competing interest.

