## Supplementary Information for "Lesion network mapping of depressive rumination identifies candidate brain stimulation targets"

### Supplementary methods

#### *fMRI data analysis*

The preprocessing of fMRI data in DIRECT was conducted using the toolbox for Data Processing & Analysis for Brain Imaging on Surface V1.8_220915 (DPABISurf)[1], which is based on fMRIPrep [2], FreeSurfer [3], ANTs [4], FSL [5], AFNI [6], SPM [7], PALM [8], dcm2niix [9], GNU Parallel [10], MATLAB R2020a (The MathWorks Inc., Natick, MA, US), Docker (https://docker.com), and DPABI V6.2_220915 [11]. The preprocessing pipeline included: 1) discarding the initial 10 time points; 2) converting data into BIDS format [12]; 3) calling fMRIPrep 20.2.1 docker; 4) conducting anatomical data preprocessing as follows: correcting the T1-weighted image for intensity nonuniformity using N4BiasFieldCorrection [13]; the corrected T1-weighted image was then skull-stripped with ANTs [4]; fast (FSL 5.0.9)[14] was used to segment brain tissue into cerebrospinal fluid (CSF), white-matter (WM) and gray-matter (GM); recon -all (FreeSurfer 6.0.1)[3] was used to reconstruct brain surfaces; ANTs-derived brain masks and FreeSurfer-derived segmentations of the cortical GM were reconciled using a custom variation of the method of Mindboggle [15]; Nonlinear registration with antsRegistration (ANTs 2.3.3) was performed for the volume-based spatial normalization to one standard space (MNI152NLin2009cAsym)[16], using brain-extracted versions of both T1w reference and the T1w template; 5) conducting functional data preprocessing as follows: generating a reference volume and its skull-stripped version using a custom methodology of fMRIPrep. The BOLD reference was then co-registered to the T1w reference using bbregister (FreeSurfer), which implements boundary-based registration [17]. Co-registration was configured with six degrees of freedom. Head-motion parameters with respect to the BOLD reference (transformation matrices and six corresponding rotation and translation parameters) were estimated before any spatiotemporal filtering using mcflirt (FSL 5.0.9). BOLD runs were slice-time corrected using 3dTshift from AFNI 20160207 (RRID:SCR_005927)[6]. The BOLD time series were resampled into standard space, generating a preprocessed BOLD run in MNI152NLin2009cAsym space; confounding time series were calculated based on the preprocessed BOLD series in the original space: framewise displacement (FD), DVARS, and three region-wise global signals (extracted from the cerebrospinal fluid (CSF), the white matter (WM), and the whole-brain masks). FD was computed using Jenkinson (relative root mean square displacement between affines) and referred to as $FD_{Jenkinson}$ [5]. 6) Nuisance regression: The Friston 24-parameter model [18] was used to regress out head motion confounds. Additionally, mean framework displacement (FD) was used to address the residual effects of motion in group analyses [5]. Other sources of spurious variance (WM and CSF signals) were also removed from the data through linear regression to reduce respiratory and cardiac effects. We performed preprocessing pipelines with and without global signal regression (GSR). Additionally, linear trends were included as a regressor to account for drifts in the blood oxygen level-dependent (BOLD) signal. vii) Finally, a bandpass temporal filter (0.01–0.1 Hz) was applied to the normalized functional images. For fMRI data in RST-Suzhou and RST-Beijing, an identical preprocessing pipeline was adopted, except for including an additional regression covariate denoting the four stimuli used in Rumination State Task (RST).

*Sample selection for the DIRECT cohort*

We selected the final sample for the image analyses according to the following criteria for DIRECT Phase II data: 1) Bad image coverage. We excluded participants with imaging data that covered less than 90 percent of the group mask. The mask denoting the analysis area was generated by extracting all brain regions that 90% of all participants' brain images covered. This discarded 15 participants. 2) Excessive head motion. Individuals with mean $FD_{Jenkinson}$ larger than 0.2 mm were excluded. Eighty-two individuals were discarded per this criterion 3) Missing demographical data. Next, we removed participants without necessary demographical information (i.e., age, sex, education, and head motion), which excluded 20 participants. This yielded a final sample containing 1574 MDD patients and 1308 healthy controls (HC).

*Harmonization of site effects*

ComBat models site-specific scaling factors by introducing terms for biological variables and scanners in a multivariate linear mixed-effects regression. Using empirical Bayesian criteria improves estimation for studies with small sample sizes. ComBat assumes that the expected FC value can be modeled as a linear combination of batch effects and biological effects. Specifically, the error term in this model is modulated by additional site-specific scaling factors. Let y denote a given functional connectivity (FC) value:

$$y = \alpha + X^T\beta + \gamma + \delta\varepsilon \quad (1)$$

where $\boldsymbol{\alpha}$ denotes the constant term; $X^T$ denotes the design matrix for covariates of interest (diagnosis, age, sex, education, head motion); $\boldsymbol{\beta}$ is a vector of regression coefficients corresponding to X; $\boldsymbol{\gamma}$ stands for additive site effects (location parameter) while $\boldsymbol{\delta}$ represents multiplicative site effects (scale parameter). Accordingly, harmonized FC values were defined as:

$$y^{ComBat} = \frac{y-\hat{\alpha}-X^T\hat{\beta}-\hat{\gamma}}{\hat{\delta}} + \alpha + X^T\hat{\beta} \quad (2)$$

where $\hat{\delta}$ and $\hat{\gamma}$ stand for the empirical Bayesian estimates of $\boldsymbol{\delta}$ and $\boldsymbol{\gamma}$, respectively. $y^{ComBat}$ was calculated with the *harmonizationLearn* function in python package *neuroHarmonize*. ComBat models biological and non-biological effects and algebraically removes the estimated additive and multiplicative batch effects.

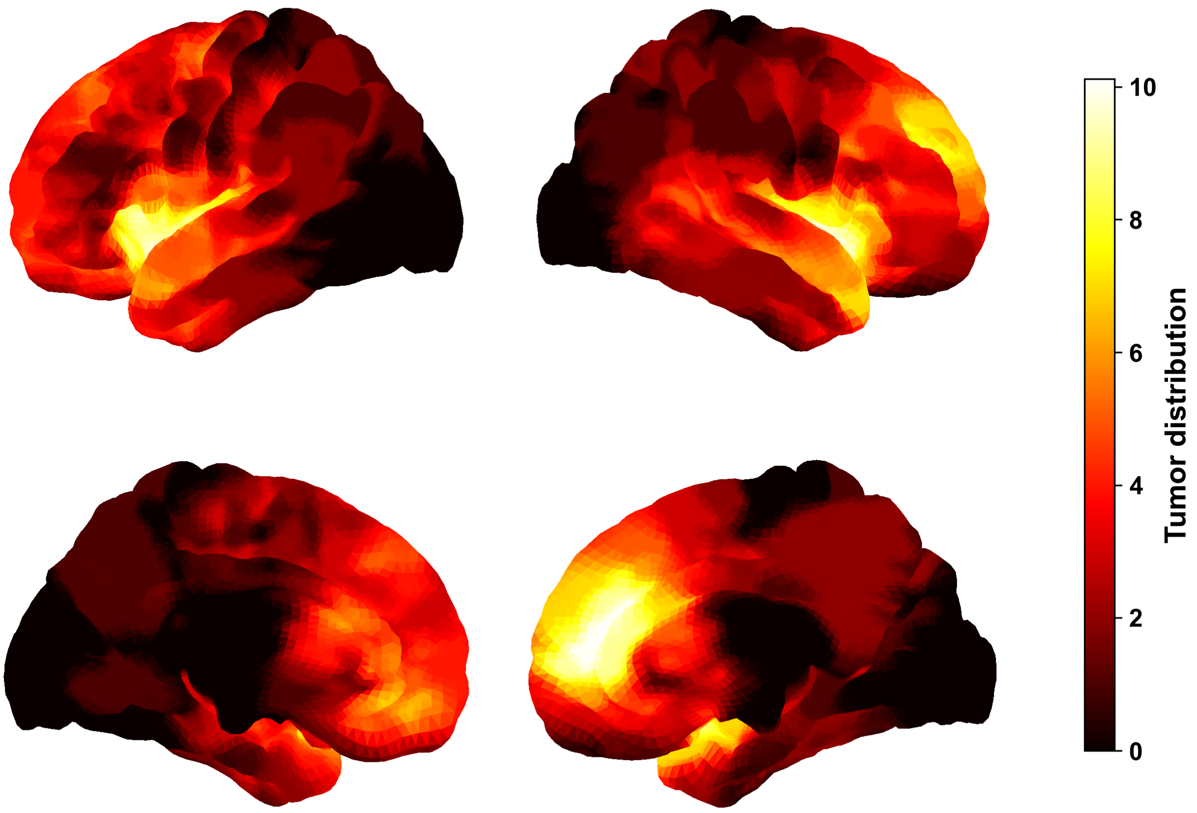


Figure 1. Spatial distribution of tumors.

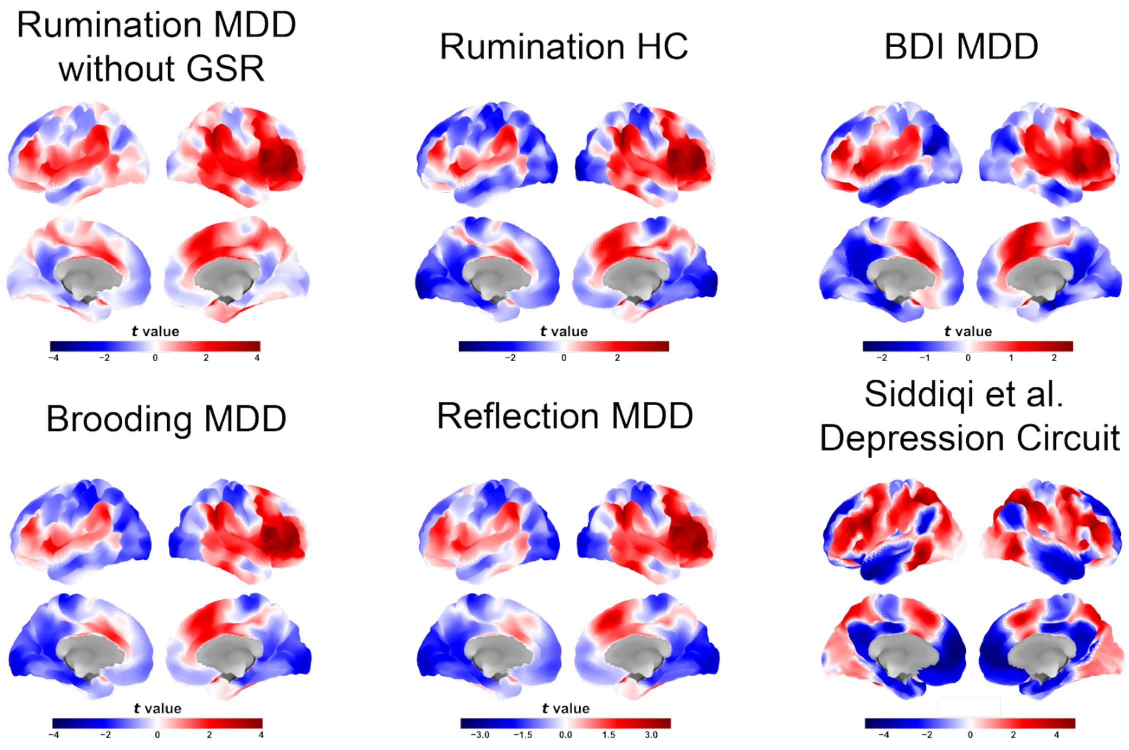


Figure S2. Comparison of rumination-related maps across preprocessing approaches, participant groups, and symptom measures. Top row, from left to right: rumination circuits in participants with MDD without GSR, rumination in HC, and the depression circuit derived using the BDI scores in participants with MDD. Bottom row, from left to right: circuits derived using the brooding and reflection subscale scores in participants with MDD, and the depression circuit reported by Siddiqi et al. [19] Abbreviations: BDI, Beck Depression Inventory; GSR, global signal regression; HC, healthy controls; MDD, major depressive disorder.

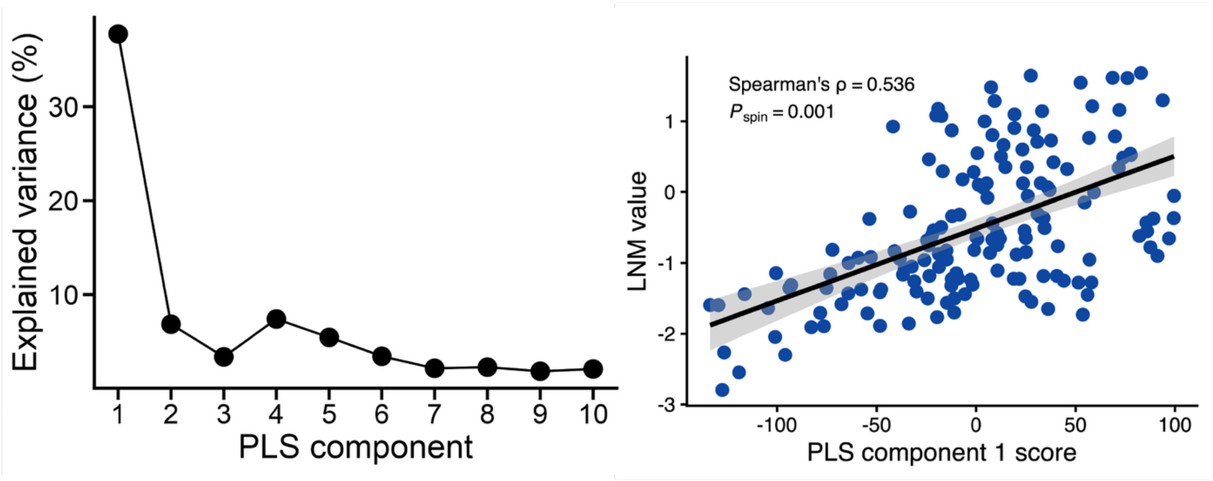


Figure S3. PLS analysis of the association between the rumination lesion network map and the gene expression profile. Left: percentage of variance explained by each of the first 10 PLS components, with the first component accounting for approximately 38%. Right: spatial association between PLS component 1 scores and LNM values (Spearman's $\rho$ = 0.536, $P_{spin}$ = 0.001). Each point represents a brain region. The solid line shows the fitted linear regression, and the shaded band indicates its confidence interval. Statistical significance was assessed using a spatial spin permutation test. Abbreviations: LNM, lesion network mapping; PLS: partial least squares.

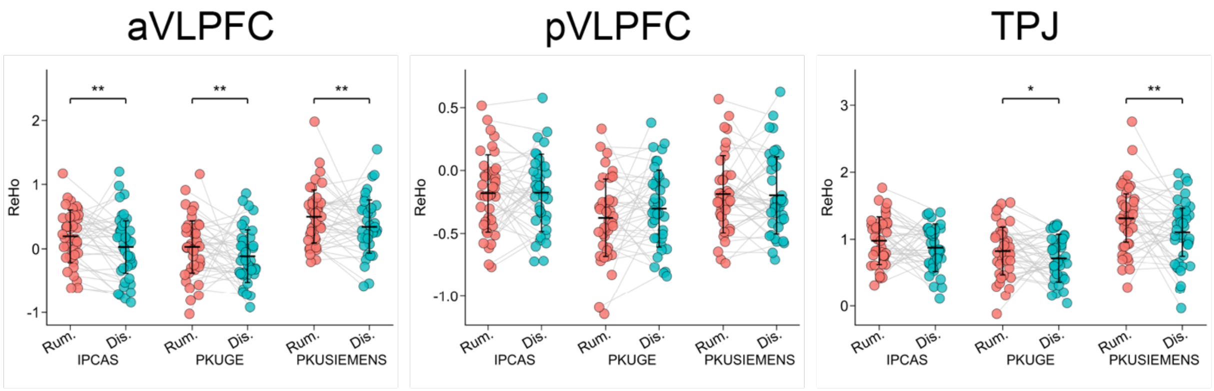


Figure S5. ReHo during rumination and distraction across three scanners in RST-Beijing dataset. Paired plots show ReHo during rumination (Rum.; red) and distraction (Dis.; blue). Each point represents one participant, and gray lines connect measurements from the same participant across conditions. Asterisks indicate significant within-dataset differences between conditions: *, $P < 0.05$ (uncorrected); **, $P < 0.05$ (corrected for multiple comparison). Abbreviations: ReHo, Regional homogeneity; TPJ, temporal parietal junction, TPL; VLPFC, ventrolateral prefrontal cortex.

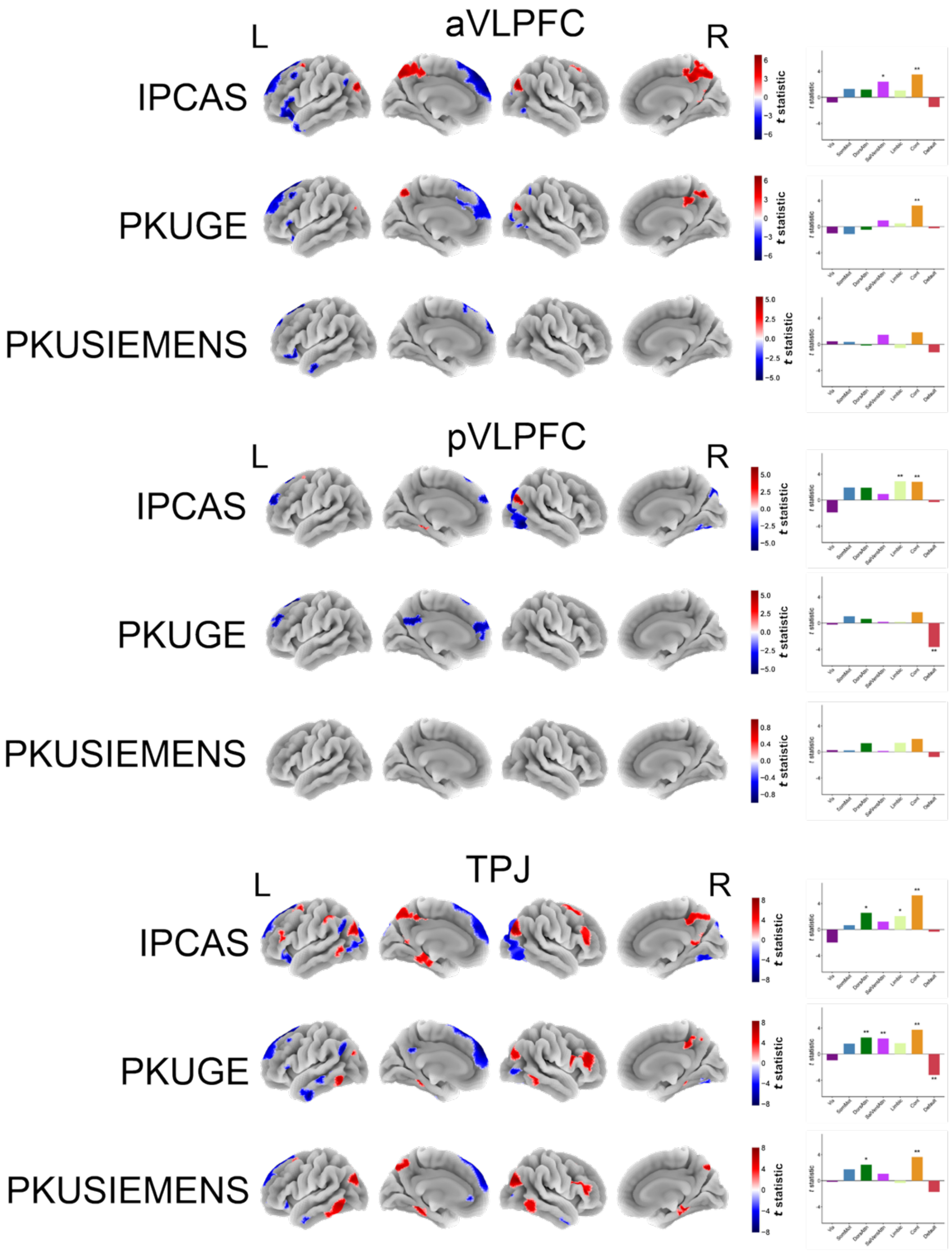


Figure S6. Functional connectivity patterns associated with candidate stimulation targets across three scanners in the RST-Beijing dataset. Bar plots summarize the corresponding results across seven functional networks, with asterisks marking statistical significance. *, $P < 0.05$ (uncorrected); **, $P < 0.05$ (corrected for multiple comparison).
